# C1q binding to surface-bound IgG is stabilized by C1r_2_s_2_ proteases

**DOI:** 10.1101/2021.02.08.430229

**Authors:** Seline A. Zwarthoff, Kevin Widmer, Annemarie Kuipers, Jürgen Strasser, Maartje Ruyken, Piet C. Aerts, Carla J.C. de Haas, Deniz Ugurlar, Gestur Vidarsson, Jos A. G. Van Strijp, Piet Gros, Paul W.H.I. Parren, Kok P.M. van Kessel, Johannes Preiner, Frank J. Beurskens, Janine Schuurman, Daniel Ricklin, Suzan H.M. Rooijakkers

**Affiliations:** Medical Microbiology, University Medical Center Utrecht, Utrecht University, Utrecht, The Netherlands; Pharmaceutical Sciences, University of Basel, Basel, Switzerland; University of Applied Sciences Upper Austria, 4020 Linz, Austria; Crystal and Structural Chemistry, Utrecht University, Utrecht, The Netherlands; Experimental Immunohematology, Sanquin Research, Amsterdam, The Netherlands; Immunohematology and Blood Transfusion, Leiden University Medical Center, Leiden, The Netherlands; Lava Therapeutics, Utrecht, The Netherlands; Genmab, Utrecht, The Netherlands

## Abstract

Complement is an important effector mechanism for antibody-mediated clearance of infections and tumor cells. Upon binding to target cells, the antibody’s constant (Fc) domain recruits complement component C1 to initiate a proteolytic cascade that generates lytic pores and stimulates phagocytosis. The C1 complex (C1qr_2_s_2_) consists of the large recognition protein C1q and a heterotetramer of proteases C1r and C1s (C1r_2_s_2_). While interactions between C1 and IgG-Fc’s are believed to be mediated by the globular heads of C1q, we here find that C1r_2_s_2_ proteases affect the capacity of C1q to form an avid complex with surface-bound IgG molecules (on various DNP-coated surfaces and pathogenic *Staphylococcus aureus*). The extent to which C1r_2_s_2_ contribute to C1q-IgG stability strongly differs between human IgG subclasses. Using antibody engineering of monoclonal IgG we reveal that hexamer-enhancing mutations improve C1q-IgG stability, both in absence and presence of C1r_2_s_2_. In addition, hexamer-enhanced IgGs targeting *S. aureus* mediate improved complement-dependent phagocytosis by human neutrophils. Altogether, these molecular insights into complement binding to surface-bound IgGs could be important for optimal design of antibody therapies.

## Introduction

Antibodies are important mediators of the human complement response, which offers critical protection against microbial infections and damaged host cells ^1^. In order to initiate a complement response, an antibody molecule first needs to bind antigens on the target cell via its antigen-binding (Fab) domains ^2–5^. Subsequently, the antibody’s constant (Fc) domain recruits the first complement protein complex, C1, to the cell surface (**SFig. 1A**). The large C1 complex (also denoted as C1qr_2_s_2_, 766 kDa) consists of the recognition protein C1q (410 kDa) and a heterotetramer of serine proteases C1r and C1s (denoted C1r_2_s_2_, 356 kDa) (**SFig. 1B**). While C1q is responsible for antibody recognition, its attached proteases C1r_2_s_2_ induce the activation of downstream enzymatic complexes, i.e. C3 convertases (C4b2b ^6^), that catalyze the covalent deposition of C3-derived molecules (e.g. C3b and its degradation product iC3b) onto the target cell surface (**SFig. 1A**) ^7,8^. C3b opsonizes the target cell surface and can induce formation of lytic pores (membrane attack complex: MAC) in the target cell membrane ^9–11^. In contrast to human cells and Gram-negative bacteria, Gram-positive bacteria are not susceptible to the MAC due to their thick cell wall ^12^. On these bacteria, efficient decoration with C3b and iC3b is essential for triggering effective phagocytic uptake of target cells via complement receptors (CR) expressed on phagocytes of which the integrin CR3 (also denoted CD11b/CD18) is considered most important ^13,14^.

In recent years, our insights into IgG-dependent complement activation have increased significantly. A combination of structural, biophysical and functional studies revealed that surface-bound IgG molecules (after Fab-mediated antigen binding) require organization into higher ordered structures, namely hexamers, to induce complement activation most effectively ^15–19^. Hexamerized IgGs are being held together by non-covalent Fc-Fc interactions and form an optimal platform for C1q docking (**SFig. 1A**). C1q has a ‘bunch of tulips’-like structure, consisting of six collagen arms that each end in a globular (gC1q) domain (**SFig. 1B**) that binds the Fc region of an IgG. As the affinity of C1q for a single IgG is very weak ^20,21^, avidity achieved through simultaneous binding of all six globular domains to six oligomerized IgG molecules is paramount for a strong response ^15,17–19^. Furthermore, it was found that IgG hexamerization could be manipulated by specific point mutations in the Fc-Fc contact region that enhance such oligomerization ^15,18,22^. While these hexamer-enhancing mutations in IgG potentiate the efficacy of MAC-dependent cytotoxicity on tumor cells and Gram-negative bacteria ^15,23^, their effect on complement-dependent phagocytosis is not known.

Because complement is an important effector mechanism to kill bacteria and tumor cells, development of complement-enhancing antibodies represents an attractive strategy for immune therapies ^1,24^. Immunotherapy based on human monoclonal antibodies is not yet available for bacterial infections ^25–28^. Such developments are mainly hampered by the fact that little is known about the basic mechanisms of complement activation on bacterial cells. For instance, we do not understand why certain antibodies induce complement activation on bacteria and others do not. In this study we set-out to investigate how antibacterial IgGs induce an effective complement response. By surprise, we noticed that C1q-IgG stability differs between human IgG subclasses. More detailed molecular investigations revealed that C1r_2_s_2_ proteases are important for generating stable C1q-IgG complexes on various target surfaces. Furthermore, we demonstrate that C1q-IgG stability is influenced by antibody oligomerization. These molecular insights into C1q binding to surface-bound IgGs may pave the way for optimal design of antibody therapies.

## Results

### IgG-mediated complement activation does not always correlate with detection of C1q

To enhance our understanding of complement activation by antibacterial IgGs, we studied complement activation by monoclonal antibodies against *Staphylococcus aureus*, an important Gram-positive pathogen and the leading cause of hospital-acquired infections. We generated IgGs against wall teichoic acid (WTA), a highly abundant and immunogenic cell wall glycopolymer that comprises 40% of the staphylococcal cell wall ^29–31^. The variable domains of anti-WTA IgG1 4497 ^31^ were cloned into HEK expression vectors encoding IgG1, IgG2, IgG3 and IgG4 Fc backbones. We included all IgG subclasses to obtain a better understanding of their variable complement effector functions ^1,32^. After confirming that all IgG subclasses bound similarly to the bacterial surface (**SFig. 2**), we examined complement activation by anti-WTA IgGs by incubating *S. aureus* with human serum as a complement source. To exclude involvement of naturally occurring anti-staphylococcal IgGs, we used serum that is depleted of natural IgG and IgM (ΔIgGΔIgM serum) ^33^. Complement activation was first quantified by measuring deposition of C3 cleavage products on the surface of *S. aureus* using flow cytometry (**Fig. 1A**). In line with recent results, IgG1 and IgG3 antibodies against WTA elicit effective C3b deposition on *S. aureus* (**Fig. 1A**) ^34^. Furthermore, IgG4 did not activate complement on *S. aureus* (**Fig. 1A**), which was expected because IgG4 has a reduced capability to interact with C1q _35–37, 38–41_.

**Figure 1.**
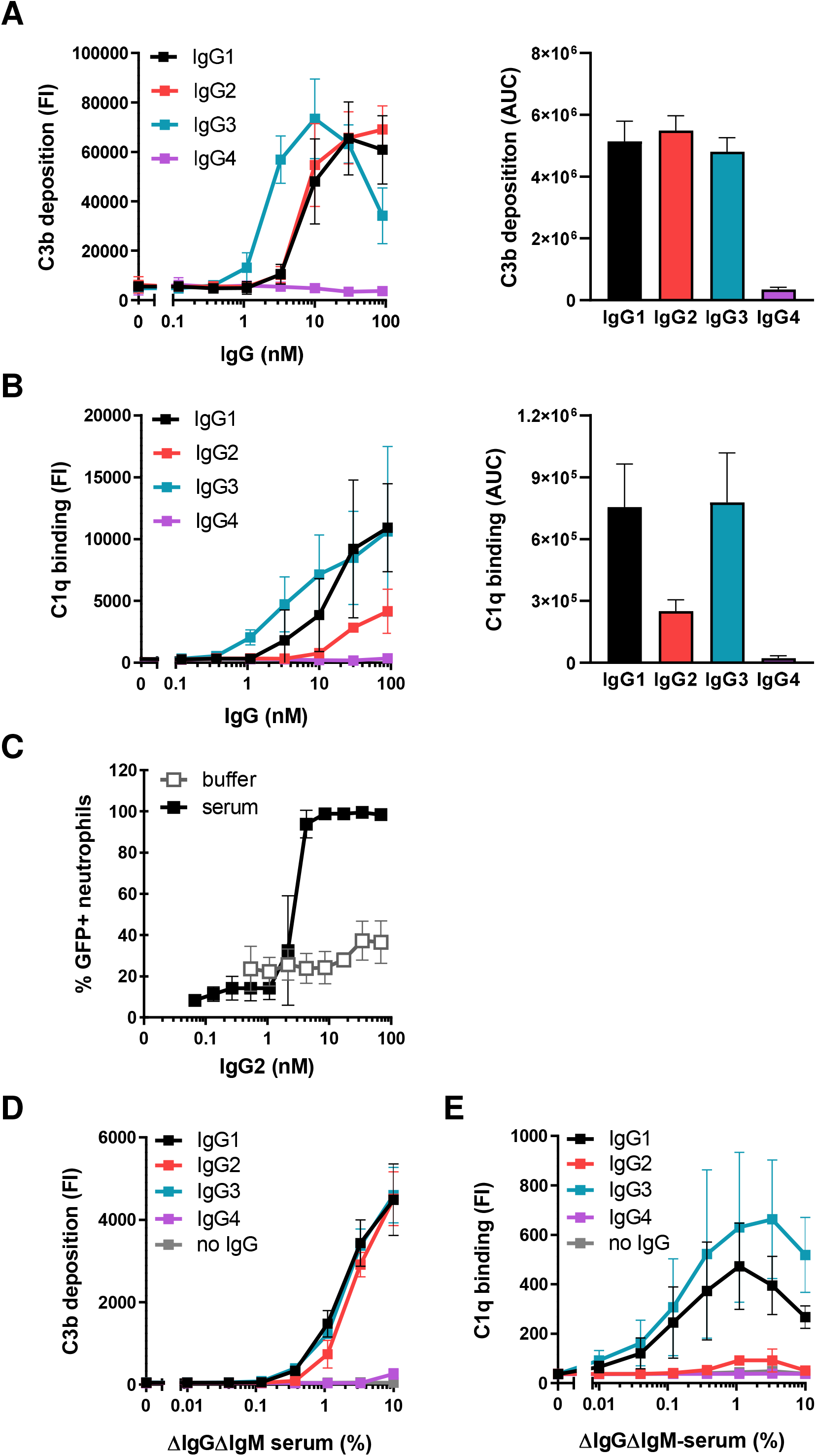
IgG-mediated complement activation does not always correlate with detection of C1q. **(A-B)** C3b deposition **(A)** and C1q binding **(B)** on anti-WTA IgG-labelled *S. aureus* Wood46 bacteria upon incubation with 5% or respectively 1% ΔIgGΔIgM serum as determined by flow cytometry. Data represent mean fluorescence intensity (FI) ± SD (left graph) or mean area under the curve (AUC) ± SD (right graph) of three independent experiments. AUC of C3b and C1q binding curves was determined after subtraction of a 2500 (C3b) or 200 (C1q) baseline, respectively. **(C)** Phagocytosis of fluorescently labelled *S. aureus* Wood46 in either RPMI buffer or 1% ΔIgGΔIgM serum supplemented with anti-WTA IgG2 and human neutrophils. Bacterial uptake was quantified by flow cytometry and displayed as the percentage of GFP-positive neutrophils. Data represent mean ± SD of three independent experiments. **(D-E)** C3b deposition **(D)** and detection of C1q **(E)** on beads coated with 1 µg/ml DNP upon incubation with ΔIgGΔIgM serum and human monoclonal anti-DNP IgG (20 nM). Deposition of C3b and C1q molecules on the beads was determined by flow cytometry. Data represent geometric mean ± SD of three independent experiments.

Our particular interest was in IgG2, which is the predominant IgG subclass against WTA in a natural human immune response ^30,42^. Although IgG2 is often described as a poor complement activator, we here observed that anti-WTA IgG2 effectively induced C3b deposition on *S. aureus*, to a level that was comparable to anti-WTA IgG1 and IgG3 (**Fig. 1A**). These data support previous studies suggesting that IgG2 can activate complement when reacting with highly dense epitopes ^42–45^.

When we took a closer look at different complement activation steps, we noticed an unexpected disparity between surface detection of C3b and C1q for IgG2. While recruitment of C1q is a prerequisite to initiate antibody-dependent complement activation, we observed that detection of C1q molecules on IgG2-coated bacteria was low compared to IgG1 and IgG3 (**Fig. 1B**). This was surprising because C3b deposition via these subclasses was similar (**Fig. 1A**). Using a monoclonal antibody that prevents C1q-IgG interactions ^46^, we showed that C3b deposition via IgG2 was driven by C1 (**SFig. 3**). Also, inhibition of C1 via a bacterial protein that blocks C1r (BBK32 ^47–49^) confirmed that C1 is required to deposit C3b onto IgG2-coated bacteria (**SFig. 3**). Furthermore, we showed that C3b molecules deposited by anti-WTA IgG2 antibodies are functional. We studied phagocytosis of bacteria by human neutrophils, the primary mechanism for elimination of *S. aureus* ^50^. Although anti-WTA IgG2 did not potently induce Fc receptor-mediated phagocytosis of *S. aureus* (as expected from the predicted low affinity of IgG2 for FcγR ^32^), we found that addition of complement strongly promoted the phagocytic uptake of IgG2-labelled *S. aureus* (**Fig. 1C**).

We wondered whether these findings could be translated to IgGs recognizing a different antigenic surface. To study this, we used an assay system in which beads are coupled with the model antigen 2,4-dinitrophenol (DNP) ^51^ and coated with human IgGs specific for DNP (**SFig. 4A-C**) ^52^. To mimic the highly abundant nature of WTA, beads were coated with saturating levels of DNP (quantified by measuring IgG binding using flow cytometry) (**SFig. 4B**). In accordance with our findings on *S. aureus*, we found that anti-DNP antibodies of the IgG1, IgG2 and IgG3 subclasses all potently induced a complement response and deposit C3b molecules onto the surface of DNP-beads (**Fig. 1D**). Again, whereas C3b opsonization could be correlated with the presence of C1q on beads coated with IgG1 and IgG3, almost no C1q could be detected on the IgG2-coated surface (**Fig. 1E)**.

In conclusion, on two independent surfaces, we showed that IgG1, IgG2 and IgG3 can all potently elicit C1-dependent deposition of C3b molecules. However, for IgG2 our data revealed an unexpected disparity between the detection of C1q and downstream deposition of C3b molecules.

### C1r_2_s_2_ proteases enhance the binding of C1q to target-bound IgG

To better understand the above findings, we more closely examined the molecular interactions between C1q and target-bound IgGs by using purified C1 complexes. We included two forms of C1q in our analyses: 1) C1q in complex with C1r_2_s_2_ proteases (denoted C1), which is representative for circulating C1 complexes in human blood ^53,54^; or 2) recognition molecule C1q without proteases (denoted C1q) (**Fig. 2A**). When we studied binding of different forms of C1q to anti-DNP antibodies on DNP-beads, we noticed a discrepancy between the binding of non-complexed C1q molecules versus C1 (**Fig. 2B, SFig. 4D**). Particularly in the case of IgG2, we observed that binding of C1 was more efficient than binding of non-complexed C1q, as quantified by flow cytometric detection of surface-bound C1q (**Fig. 2B**). Western blotting was used to confirm that the different levels of C1q detected in flow cytometry actually represent different quantities of surface-bound C1q (**SFig. 4E**). When C1 complexes were dissociated by EDTA, which disrupts the calcium-dependent attachment of C1r_2_s_2_ to C1q ^55,56^, binding was similar to C1q alone (**Fig. 2B**). On IgG1-coated beads, we observed that C1r_2_s_2_ proteases had subtle effect on binding of C1q (**Fig. 2B**). Furthermore, C1r_2_s_2_ proteases had a very limited effect on the binding of C1q to IgG3 (**Fig. 2B)**. Similar to these results on beads, we observed that EDTA reduced binding of purified C1 to *S. aureus* labelled with IgG1 and IgG2, while binding to IgG3 was much less affected (**SFig. 5**). Altogether, these studies suggest that attached C1r_2_s_2_ proteases affect C1q-IgG interactions in a subclass-dependent manner. This finding is unexpected when considering that the direct interactions between C1 and IgG were shown to solely depend on the gC1q domains ^16^.

**Figure 2.**
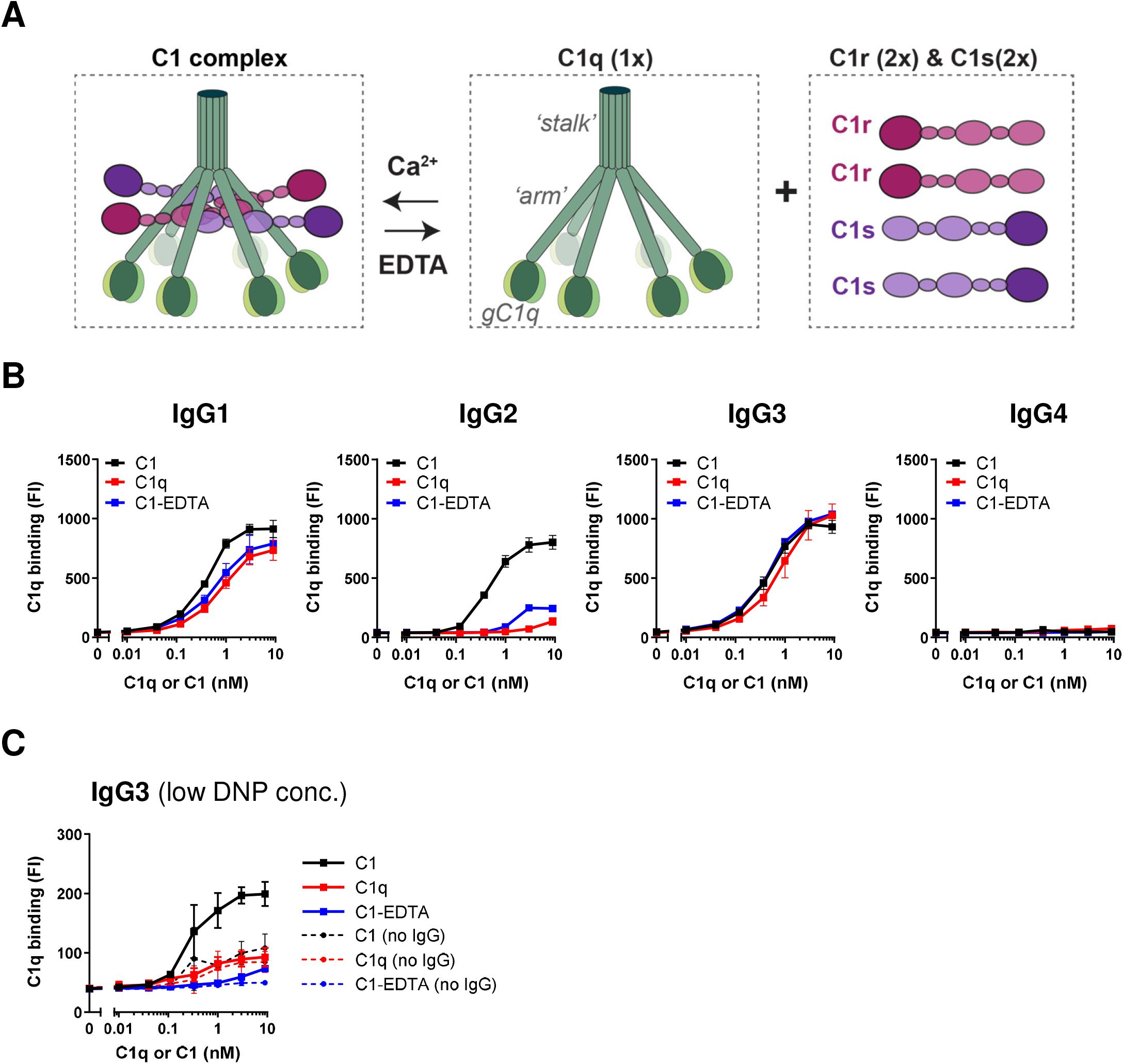
C1r_2_s_2_ proteases enhance the binding of C1q to target-bound IgG. **(A)** Schematic presentation of C1 consisting of the recognition molecule C1q in complex with a tetramer of proteases C1r and C1s (C1r_2_s_2_). C1q consists of six polypeptide chains that come together in an N-terminal stalk. At their C-terminus, all six chains end in a globular domain (gC1q) that recognizes Fc domains of IgM and clustered IgGs. The C1r_2_s_2_ tetramer associates to the collagen arms of C1q via Ca^2+^-dependent interactions ^56^. The proteases dissociate from C1q in the presence of EDTA, a calcium-chelator. **(B)** Binding of different forms of purified C1 to DNP-beads (1 µg/ml DNP) coated with 20 nM anti-DNP IgG1-4. ‘C1’ indicates the fully assembled C1 complex (= C1qr_2_s_2_), ‘C1q’ is the recognition molecule C1q only, ‘C1-EDTA’ sample consists of C1q, C1r and C1s, but the proteases are not attached to C1q (because 10 mM EDTA disrupts the Ca^2+^-dependent association between proteases and C1q). **(C)** Binding of different forms of purified C1 to IgG3-labelled beads coated with 0.003 µg/ml (∼300-fold lower than in **(B)**). The dotted lines show aspecific binding of the C1q molecules in absence of IgG3. **(B-C)** Bound C1q was detected by polyclonal anti-C1q antibodies and flow cytometry. Data represent mean ± SD of three independent experiments.

Since the surface density of antigens was earlier proposed to be a critical parameter for IgG-mediated complement activation ^57–59^, we wondered whether C1r_2_s_2_ proteases also affect C1q-IgG interactions at lower antigen concentrations. In the bead system, we lowered the DNP concentration by ∼300-fold. At this lower DNP concentrations, we found that IgG3 was the most potent subclass triggering complement activation in human serum (**SFig. 6**) and that C3b deposition by IgG1 and IgG2 was inefficient. This corresponds with the idea that IgG3 more potently drives complement activation on low abundant antigens (presumably because it has a longer hinge region ^21,57,58,60^). Upon studying binding of purified C1 complexes, we observed that C1r_2_s_2_ proteases can also affect C1q binding to IgG3 on beads with a lower DNP concentration. First, we observed that C1 bound more efficiently to IgG3 than non-complexed C1q (**Fig. 2C**). Also, disruption of C1 with EDTA caused an almost complete reduction of C1q binding to IgG3 (**Fig. 2C**).

Altogether this suggests that C1r_2_s_2_ proteases can enhance binding of C1q to all IgG subclasses. However, the extent to which C1r_2_s_2_ proteases contribute to C1q binding depends both on the IgG subclass and antigen concentration.

### C1r_2_s_2_ proteases enhance the stability of surface-bound C1q-IgG complexes

To corroborate these results, we employed surface plasmon resonance (SPR) as an orthogonal technique to confirm the binding profiles and obtain dynamic information about the formation and stability of C1q-IgG complexes (**Fig. 3A, SFig. 7**). Using a flat DNP-labelled surface, prepared by coupling DNP-PEG-NHS to an activated carboxyl sensor chip (**SFig. 7A**), monoclonal anti-DNP antibodies could be captured at high density and stability (**SFig. 7B**). No binding was observed on a MeO-PEG-NHS surface that was used as a reference (**SFig. 7C**). By injecting either C1q or C1 for 60 seconds, the assembly (during injection) and stability of C1q-IgG/C1-IgG complexes (during the dissociation period) could be monitored over time. SPR analysis indeed confirmed that C1r_2_s_2_ proteases affect the stability of C1q-IgG complexes, especially for IgG1 and IgG2 (**Fig. 3A**). Whereas both C1q and C1 bound to the IgG1- and IgG2-coated sensor chips during protein injection, C1 dissociated much slower than C1q after the injection was stopped, which suggests that C1 forms more stable interactions with the antibody-coated surface. Additionally, we observed a weaker association of C1q in absence of C1r_2_s_2_ to IgG2-coated chips during the injection (**Fig. 3A**). In line with our studies on IgG3-labelled beads, C1r_2_s_2_ proteases did not notably affect C1q binding to IgG3-coated chips. However, SPR experiments showed that C1r_2_s_2_ proteases had a more pronounced impact on binding of C1q when the IgG3 capturing concentration was lowered 5-fold (**SFig. 7D**). As expected, neither C1q nor C1 showed detectable binding to IgG4.

**Figure 3:**
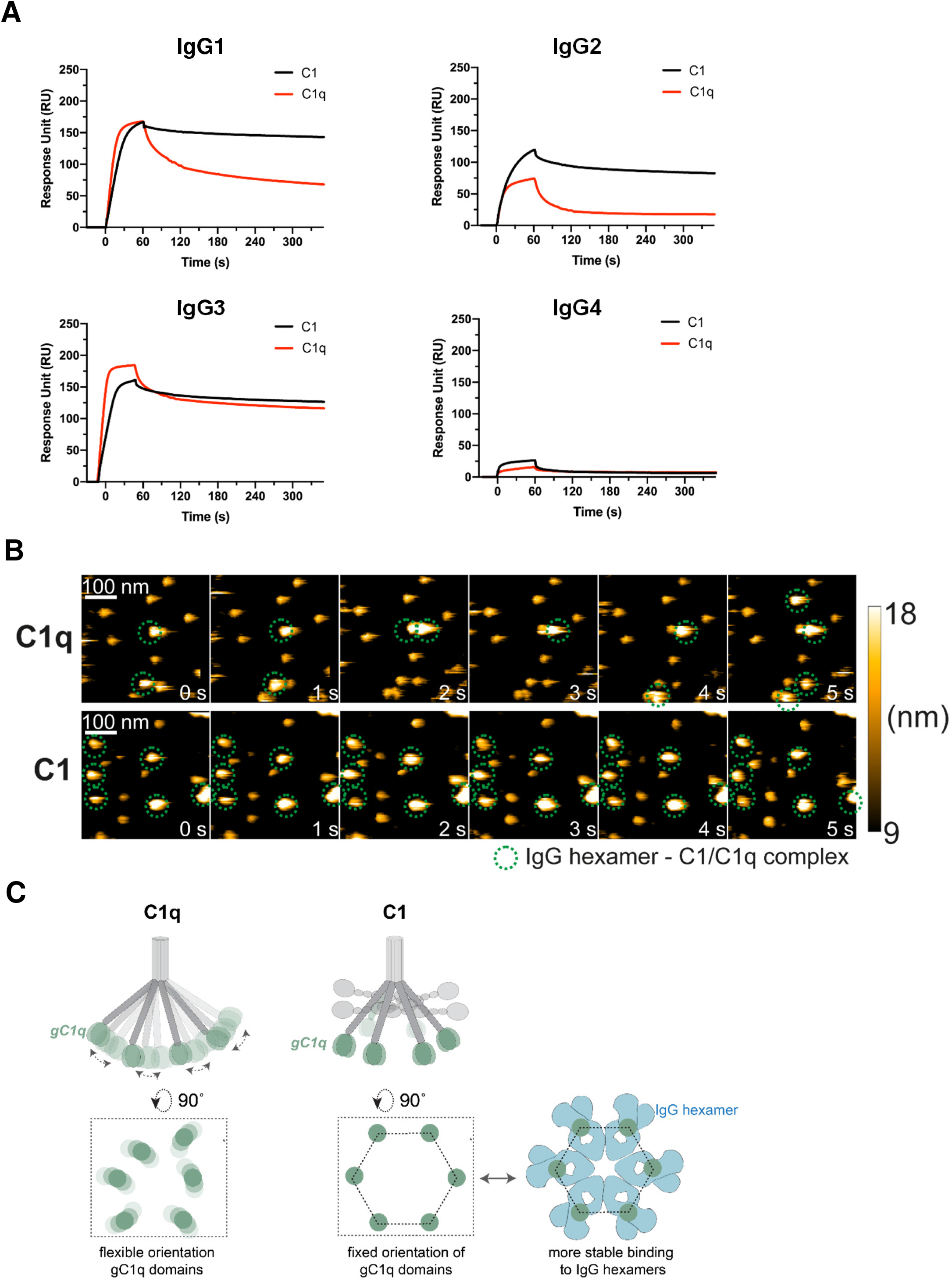
C1r_2_s_2_ proteases enhance the stability of surface-bound C1q-IgG complexes. **(A)** SPR experiment showing binding of purified C1 or C1q to sensor chips with immobilized DNP coated with 20 nM anti-DNP IgG. C1 or C1q was injected for 60 seconds to allow association, after which the injection was stopped and dissociation was monitored. Representative of two independent experiments. SPR responses were normalized to account for the molecular weight difference between C1 (766 kDa) and C1q (410 kDa). RU, response units. **(B)** HS-AFM image sequence of anti-DNP IgG1-RGY in complex with C1q in the presence of C1q in solution (upper panel; taken from **Movie S1**) and C1 in the absence of C1 in solution (lower panel; taken from **Movie S2**). The depicted height scale is relative to the membrane surface. The heights of the respective complexes were 12.2 ± 0.6 nm (anti-DNP IgG1-RGY) ^17^, 20.2 ± 1.5 nm (anti-DNP IgG1-RGY + C1q), and 18.5 ± 1.7 nm (anti-DNP IgG1-RGY + C1). **(C)** gC1q domains mediate binding to IgG. In C1q, the gC1q domains are flexible. We hypothesize that in C1 the associated C1r_2_s_2_ proteases fix the collagen arms and thereby orient the gC1q domains in a hexagon-like platform that favors binding to (hexameric) IgG clusters.

To directly visualize C1q-IgG complexes, we performed high-speed atomic force microscopy (HS-AFM) experiments. To enable reliable identification of C1q-IgG complexes in HS-AFM, we employed the anti-DNP triple mutant (IgG1-E345R, E430G, and S440Y, denoted IgG1-RGY), which was shown to efficiently associate into IgG1 hexamers in solution ^19,61^. DNP-labelled supported lipid bilayers were therefore pre-incubated with anti-DNP IgG1-RGY ^17^. Then, C1q or C1 was added and the resulting complexes where visualized via HS-AFM (**Fig. 3B, Movies S1** and **S2**). While the complexes of C1 that bound to the IgG1-RGY hexamers were not significantly disturbed by the minimal forces exerted by the HS-AFM tip, C1q alone was frequently removed from the IgG1-RGY hexamers as a result of the tip-sample interaction under the same experimental settings.

Altogether, the above data suggest that attached C1r_2_s_2_ proteases improve the stability of C1q-IgG complexes on target surfaces. We propose that C1r_2_s_2_ proteases affect the conformation of C1q in a way that facilitates stable docking to surface-bound IgGs. Earlier studies reported that the solution structure of C1q shows a high degree of flexibility ^62–64^. While the six collagen arms firmly bundle via disulphide bonds in the upper stalk, such interactions are lacking below the stalk and enable a rather flexible arrangement of the gC1q domains (**Fig. 3C**). A recent *in situ* 3D structure of C1qr_2_s_2_, bound to surface-antigen–bound IgM, showed that the C1r_2_s_2_ proteases are packed inside the C1q molecule as two antiparallel C1rs heterodimers ^65^. Each C1rs dimer binds three of the six C1q collagen helices, which limits the flexibility of the collagen arms and results in a near-hexagonal arrangement of the six gC1q domains. Based on recent observations that IgG hexamers are the ideal docking platform for C1q ^15,17^, we propose that this hexagon-like arrangement of the six gC1q domains induced by the attachment of C1r_2_s_2_ to C1q favors multivalent, high-avidity binding of C1q with clustered IgG (**Fig. 3C**).

### Removal of C1r_2_s_2_ proteases from surface-bound C1-IgG complexes can result in C1q dissociation

Next, we wondered how the observed differences in stability between C1-IgG and C1q-IgG could explain why we observed a discrepancy between C1q and C3b detection on IgG2-coated surfaces in serum (**Fig. 1**). Because C1q in human serum circulates as a complex with C1r_2_s_2_ proteases, we assume that IgG2-coated *S. aureus* and DNP-beads (1 µg/ml DNP) can recruit C1 and activate complement. However, since human serum contains an inhibitor that removes C1r_2_s_2_ proteases from C1q, we hypothesized that subsequent removal of proteases results in dissociation of C1q-IgG complexes. To test this, we incubated IgG-covered beads with purified C1 and, after washing, incubated the C1-bound beads with human C1-esterase inhibitor (C1-INH) ^66^. C1-INH is a human serpin that inactivates the proteases by forming a covalent bond with the catalytic site of both C1r and C1s. As expected, incubation of C1-IgG complexes on beads with C1-INH led to removal of C1r_2_s_2_ proteases from C1q on IgG1-, IgG2- and IgG3-covered beads as evidenced by the detection of C1-INH-C1r and C1-INH-C1s complexes in the sample supernatant using western blotting (**SFig. 8**). In line with our hypothesis, we found that the dissociation of C1r_2_s_2_ by C1-INH had differential effects on the stability of the bead-bound C1q-IgG complexes (**Fig. 4A**). Whereas dissociation of C1r_2_s_2_ proteases by C1-INH did not affect C1q-IgG3 complexes, it caused a strong (80%) reduction of C1q binding to IgG2-beads and a 10% reduction on IgG1-beads (**Fig. 4A**). Similar results were obtained when C1r_2_s_2_ proteases were removed from C1-IgG complexes using EDTA (**Fig. 4A**).

**Figure 4:**
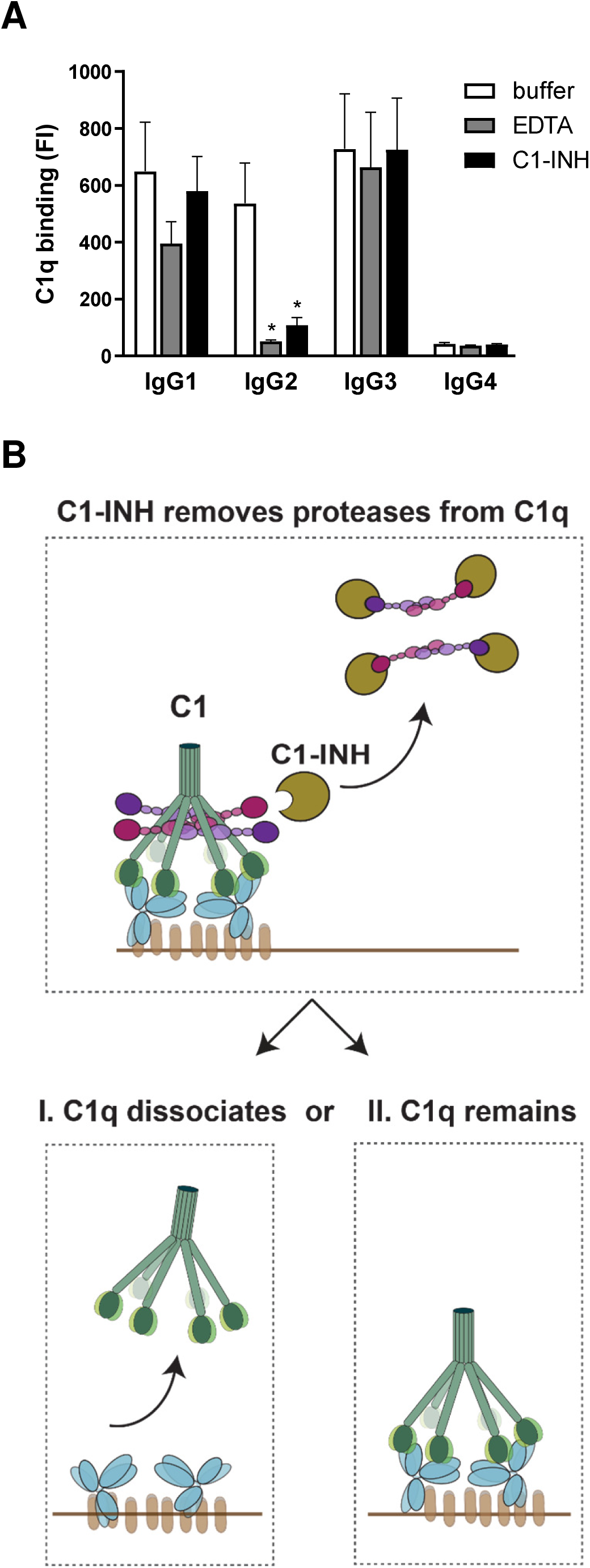
Removal of C1r_2_s_2_ proteases from surface-bound C1-IgG complexes can result in C1q dissociation. **(A)** Detection of C1q on IgG-coated DNP-beads that were first labelled with purified C1 and subsequently incubated with 10 mM EDTA or 200 nM C1-INH to remove C1r and C1s proteases. Data represent geometric mean ± SD of three independent experiments. Unpaired t test (buffer vs. EDTA; buffer vs. C1-INH); \**P* < 0.05, all other conditions not significant. **(B)** Schematic cartoon of our hypothesis that C1r_2_s_2_ dissociation by C1-INH can result in C1q dislodgement depending on the stability of the C1q-IgG complex. Our data suggest that removal of the C1r and C1s proteases from surface-bound C1 by C1-INH can result in two situations: **(I)** C1q dissociates from the surface-bound IgGs in the case the remaining C1q-IgG complexes are unstable, e.g. for C1q-IgG2 complexes; or **(II)** C1q remains bound since it has formed a stable interaction with the surface-bound IgGs.

Altogether, these data demonstrate that C1-INH can dislodge C1q from IgG-coated surfaces by removing C1r_2_s_2_ from surface-bound C1-IgG complexes. While C1-INH binds and removes C1r_2_s_2_ regardless of the IgG subclass, subsequent C1q dissociation depends on the stability of C1q-IgG complexes (**Fig. 4B**). It seems likely that the dislodgement of C1q as result of C1r_2_s_2_ dissociation explains why we could not detect C1q on IgG2-beads in a serum environment (**Fig. 1E**). The observed deposition of C3b molecules on IgG2-beads in serum (**Fig. 1D**) furthermore suggests that C1 inhibitory mechanisms lag behind on C1-mediated cleavage of complement proteins.

### Enhanced IgG oligomerization stabilizes C1q-IgG interactions

Finally, we determined how C1q-IgG interactions are influenced by IgG oligomerization. Recent studies demonstrated that oligomerization of target-bound IgG into hexamers can be enhanced by specific point mutations that strengthen Fc-Fc contacts ^15,18^. In contrast to the RGY mutation used above, these so-called HexaBody® mutations do not enhance hexamer formation in solution but specifically enhance hexamerization on target surfaces. Here, we modified anti-DNP IgGs using hexabody mutations E430G or E345K ^18^ and verified that these mutations did not affect IgG binding to beads (**SFig. 9A-B**). Upon studying interactions with purified forms of C1/C1q, we observed that both Fc mutations strongly improved binding of non-complexed C1q (i.e. C1q or C1-EDTA) to IgG1 and IgG2 on DNP-beads (**Fig. 5A, SFig. 9C**). For IgG3, hexabody mutations did not affect binding of non-complexed C1q (**Fig. 5A, SFig. 9C)**. SPR experiments corroborated the results in the bead assay, showing that both the E430G and E345K mutation increase complex stability of IgG1 and IgG2 with C1q and have little effect on the association of C1q with IgG3 (**SFig. 10**). Similar results were obtained for *S. aureus*, where introduction of E430G into anti-WTA IgG, strongly enhanced binding of non-complexed C1q to IgG1 and IgG2, while not affecting IgG3 (**SFig. 11A-B)**. Interestingly, we observed that hexabody mutations did not have a strong impact on the binding of fully assembled C1 to IgG1 and IgG2 on DNP-coated beads (**Fig. 5B**) or SPR chips (**SFig. 10**). On *S. aureus* we observed that the E430G mutation induced a subtle improvement of C1 binding (**SFig. 11C**). These data suggest that unstable C1q-IgG interactions between non-complexed C1q and IgG can be overcome by promoting formation of high avidity multimeric IgG platforms on the surface. In situations where C1q-IgG complexes are already stable (e.g. for C1 binding to IgG1, IgG2 or IgG3 or C1q binding to IgG3), hexabody mutations do not or only slightly improve C1q-IgG binding. The observation that Fc mutation E430G enhances C1q binding to IgG1 and IgG2 in human ΔIgGΔIgM serum (both on beads (**Fig. 5C**) and *S. aureus* (**SFig. 12**)) suggests that hexabody mutations also improve the ability of IgGs to retain C1q on target surfaces in a serum environment.

**Figure 5:**
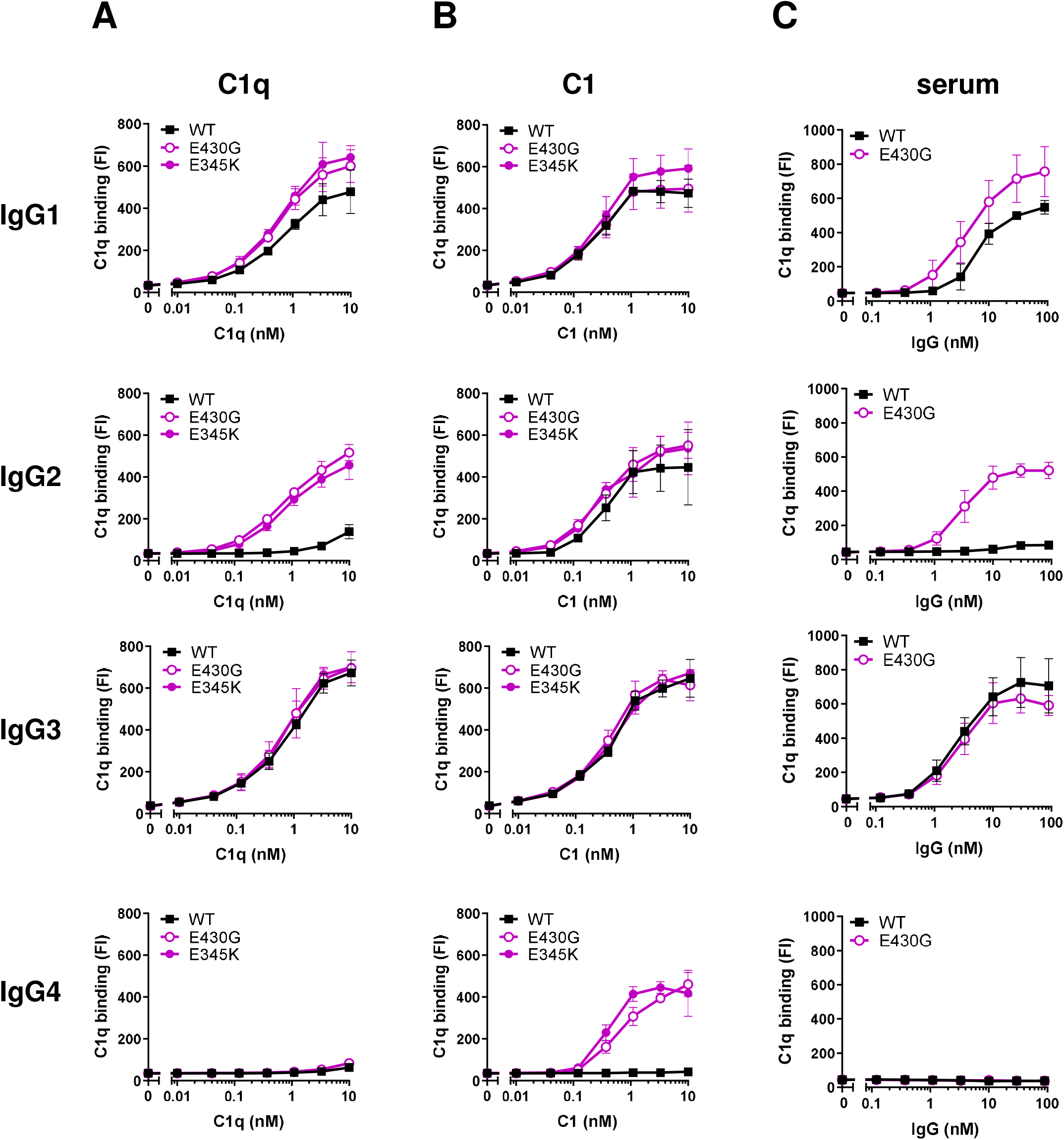
Enhanced IgG oligomerization stabilizes C1q-IgG interactions. **(A-B)** Binding of purified C1q **(A)** or C1 **(B)** to DNP-beads (1 µg/ml DNP) labelled with 20 nM anti-DNP IgG, either wild-type or containing hexamer-enhancing mutations E430G or E345K. **(C)** Detection of C1q on DNP-beads (1 µg/ml DNP) after incubation with 1% ΔIgGΔIgM serum supplemented with wild-type or mutated (E430G) anti-DNP IgG. **(A-C)** C1q was detected by flow cytometry. Data represent geometric mean ± SD of three independent experiments.

We also examined whether enhanced hexamerization could affect complement binding and activation via IgG4, the IgG subclass that is considered incapable of reacting with C1q ^39^. Previous studies showed that the Ser at position 331 in the heavy chain of IgG4 is critical for determining its inability to bind C1q ^38,67^. Structural modelling has furthermore shown that the two Fab arms of IgG4 obstruct its C1q binding site and that the short IgG4 hinge region allows only limited flexibility ^35–37^. As the IgG4 used in our study also contains the essential Ser331 residue, we expected no binding to C1q or C1. However, when we introduced the E430G or E345K mutation in anti-DNP IgG4, we observed dose-dependent binding of fully assembled C1, but not C1q (**Fig. 5A-B, SFig. 9C**). Also on *S. aureus* (**SFig. 11**) and in SPR assays (**SFig. 10**), we observed that enhancing Fc-Fc interactions in IgG4 enabled binding to the fully assembled C1, and negligible binding to non-complexed C1q. On *S. aureus* in human serum, we could detect only low levels of C1q when introducing the hexabody mutation E430G in IgG4 (**SFig. 12B**). This observation is not unexpected considering our earlier hypothesis that C1r_2_s_2_ removal by C1-INH results in dislodgement of C1q when C1q-IgG interactions are unstable. Taken together, these data suggest that the poor interaction of surface-bound IgG4 with C1q can be overcome in the presence of both hexabody mutations and C1r_2_s_2_ proteases which confirms a role for the proteases in C1q binding.

### Introduction of hexamer-enhancing mutation E430G in anti-WTA enhance complement-dependent phagocytosis of S. aureus

Having demonstrated that enhanced IgG hexamerization improves the stability of C1q-IgG complexes on *S. aureus*, we wondered whether hexabody mutations also have an impact on the downstream complement effector mechanisms. In previous studies, we and others have shown that phagocytosis of *S. aureus*, which is eminent for human immune defense against these bacteria ^68^, critically depends on the labelling of *S. aureus* with complement-derived opsonins _69–71_. While antibacterial IgGs may trigger phagocytosis of *S. aureus* via Fc receptors, additional opsonization of the bacterial surface with C3b and iC3b enhances the efficacy of particle uptake via engagement of complement receptors. To study the effect of hexamer-enhancing mutations on phagocytosis, we first determined the efficiency by which anti-WTA mutant IgGs triggered covalent deposition of opsonic C3b on *S. aureus* (**Fig. 6A**). Upon incubation of *S. aureus* with anti-WTA IgGs and ΔIgGΔIgM serum, we observed that the E430G mutation enhanced C3b deposition mediated by IgG1 and IgG2, but not IgG3 (**Fig. 6A**). Next, we determined whether enhanced IgG oligomerization on *S. aureus* improved the phagocytosis of bacteria by human neutrophils in serum (**Fig. 6B**). In full correspondence with the observed improvements at the level of C1q (**SFig. 12B**) and C3b (**Fig. 6A**), we observed that introduction of the E430G mutation in anti-WTA IgG1 and IgG2, but not IgG3, enhanced the phagocytic uptake of fluorescent *S. aureus* in human serum. Consistent with the finding that anti-WTA IgG4-E430G antibodies can induce C3b deposition, we observed that this antibody could induce phagocytosis of *S. aureus* (**Fig. 6B**). As expected, the E430G mutation did not affect the IgG-dependent phagocytosis via Fc receptors in the absence of complement (**SFig. 13**). Altogether these data demonstrate that Fc-engineering of anti-*S. aureus* IgGs can be a useful strategy to improve a complement response against this important pathogen and subsequent uptake by phagocytes.

**Figure 6:**
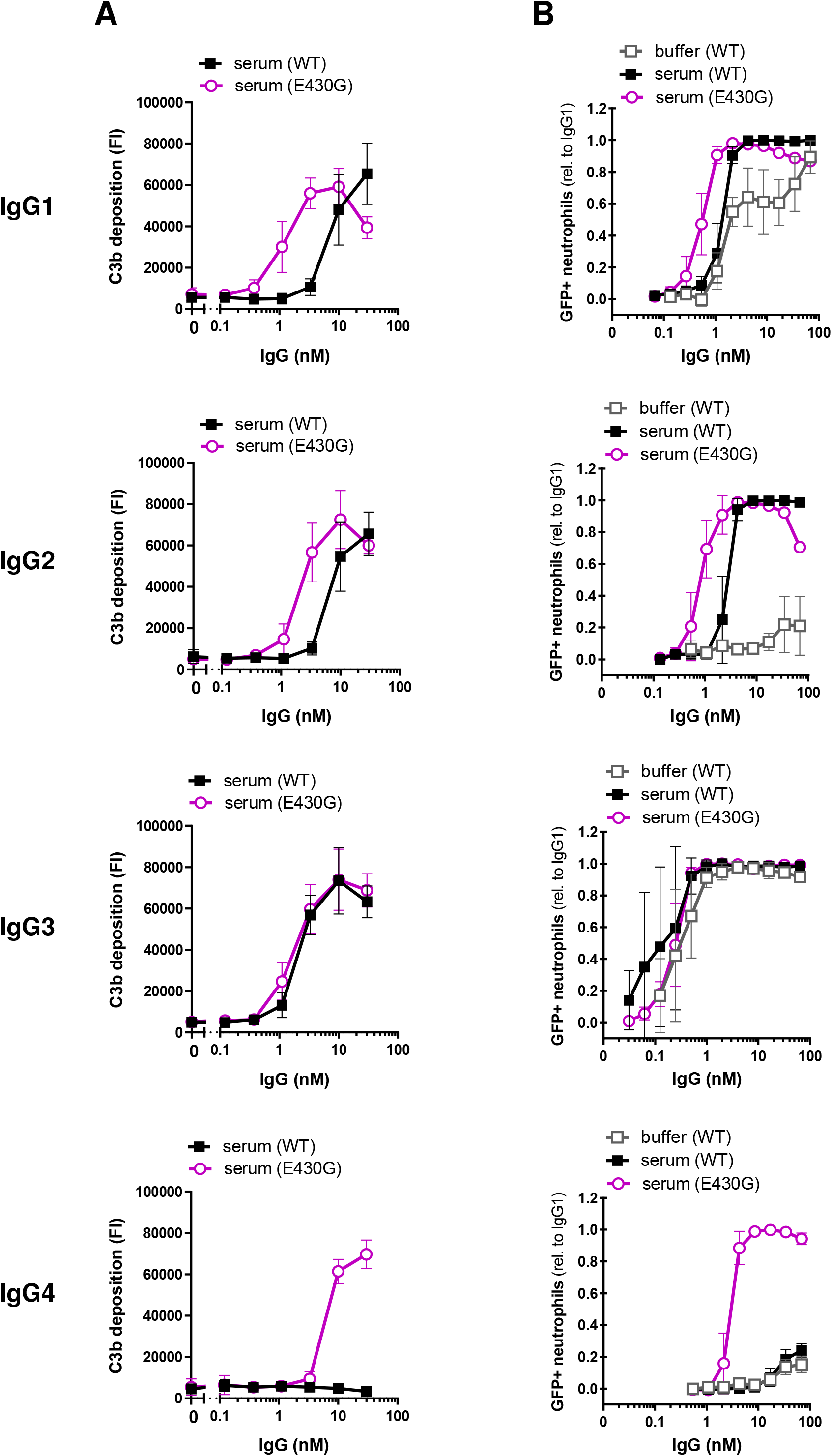
Introduction of hexamer-enhancing mutation E430G in anti-WTA can enhance complement-dependent phagocytosis of *S. aureus*. **(A)** C3b deposition on *S. aureus* Wood46 after incubation of bacteria in 5% ΔIgGΔIgM serum supplemented with anti-WTA IgG (wild-type or E430G mutant). Data represent mean ± SD of three independent experiments. **(B)** Phagocytosis in the absence and presence of complement. Phagocytosis of fluorescently labelled *S. aureus* Wood46 in either RPMI buffer or 1% ΔIgGΔIgM serum supplemented with anti-WTA IgG (wild-type or E430G mutant) and human neutrophils. Bacterial uptake was quantified by flow cytometry and displayed as the number of GFP-positive neutrophils relative to IgG1 wild-type. Data represent relative mean ± SD of three independent experiments. Phagocytosis data shown for wild-type IgG2 (buffer and serum condition) are identical to the data shown in **Fig. 1C**.

## Discussion

The classical pathway of complement activation is a major contributor to pathophysiological processes in our body, including infection, inflammation, autoimmunity and transplant rejection. Here we demonstrate in great detail how the initiating step of this pathway, namely the binding of the large C1 complex to the surface of IgG-labelled target cells, is influenced by its attached proteases C1r_2_s_2_ and antibody oligomerization. Using well-defined monoclonal human antibodies and highly purified model systems to study C1q-IgG interactions on target surfaces, we here demonstrate that C1r_2_s_2_ proteases affect the capacity of C1q to form an avid complex with surface-bound IgG molecules. Although biophysical studies demonstrated that C1r_2_s_2_ proteases modify the highly flexible structure of C1q in solution ^62–64^, it was not known how this would affect C1q binding to surface-bound IgG. Okada *et al*. (1985) proposed a potential role for C1r_2_s_2_ proteases in C1q binding to polyclonal rabbit IgG, but the mechanism and relevance for human IgG remained unclear ^72^. Since our data now show that associated C1r_2_s_2_ are important for generating stable C1q-IgG complexes on surfaces, we propose that the associated proteases limit the conformational flexibility of the C1q arms and stabilize a spatial near-hexagonal arrangement of the six gC1q domains that facilitates multivalent binding of C1q to antibody clusters on the surface (**Fig. 3C**).

Understanding how C1q molecules form a stable interaction with surface-bound IgGs is crucial for future design of therapeutic antibodies. Antibodies against bacteria and tumor cells cannot directly neutralize an infection but require activation of the immune system. For Gram-positive bacteria, the functionality of antibodies in human immune protection critically depends on the antibody’s capacity to induce phagocytic killing ^68,73^, either directly or via complement. Here we show that Fc-engineering could be a useful strategy to combat Gram-positive bacteria, as both enhancement of C1q and C1 binding to *S. aureus* can improve the opsonisation and phagocytosis of these bacteria in human serum. Recent studies on tumor cells and *Neisseria gonorrhoeae* have already shown that enhanced hexamerisation of IgG by Fc-engineering can potentiate the capacity of monoclonal antibodies to induce complement-mediated lysis via MAC pores ^15,18,19,23^. Here, we demonstrate for the first time that the same mutations can be used to improve complement-dependent phagocytosis. Furthermore, since complement activation also appears crucial to induce protective antibodies against viral infections such as HIV or SARS-CoV2 ^74,75^, designing effective complement-triggering antibodies may also be valuable for therapeutic development of anti-viral antibodies ^24^. In addition to its role in complement activation, increasing evidence suggests that C1q also enhances the efficacy of antibodies in the absence of other complement proteins. On tumor cells, it was shown that C1q can act as a structural component in potentiating the efficacy of anti-tumor antibodies by inducing outside-in signaling of death receptors on tumor cells ^76^. Furthermore, C1q directly enhances the neutralizing activity of anti-viral antibodies *in vitro* and *in vivo* ^77^.

Our study highlights that the IgG-Fc domain has a large impact on the antibody’s capacity to form stable complexes with C1q. Detailed comparison of various monoclonal antibodies indicate that the difference in stability between C1-IgG and C1q-IgG is most apparent under conditions where IgG oligomerization is less favorable. Based on our results with hexamer-enhancing mutations, we think that effective IgG oligomerization can overcome the need for proteases to establish stable C1q-IgG complexes, likely because six antibodies can also form a stable, high avidity complex with six C1q arms. Furthermore, hexamer-enhancing mutations in IgG2 and IgG1 also increased C1 binding to *S. aureus*, indicating that even when C1r_2_s_2_ is associated, efficient IgG oligomerization further stabilizes C1-IgG complexes. Interestingly, the finding that hexamer-enhanced IgG4 enabled binding to fully assembled C1, but less strongly to non-complexed C1q, could suggest that IgG4-E430G does not efficiently form hexamers. In all, we hypothesize that differences between human IgG subclasses to bind C1q (in the absence of proteases) could be linked to their ability to form hexameric platforms. However, also the binding affinity of IgG subclasses for gC1q may play a role. It remains to be determined whether our experiments on beads and Gram-positive bacteria (both rigid surfaces) can be translated to highly fluidic eukaryotic or Gram-negative membranes.

Our data also shed light on the mechanisms by which C1-INH inactivates C1, which are not yet fully defined. In 1998, Chen and Boackle already showed that purified C1-INH, besides removing C1r_2_s_2_, can dislodge C1q molecules from an IgG-coated surface. Although they used pooled human IgG in their C1-INH studies, they suggested that the IgG subclasses might be differentially affected by C1-INH ^78,79^. Our data now provide a better insight into the subclass-specific effects of C1-INH. They show that C1-INH inhibits the C1r_2_s_2_ proteases in surface-bound C1-IgG independent of the IgG subclass, but that it mediates C1q dissociation only in conditions where C1q-IgG complexes are not stable. This suggests that protease dissociation by C1-INH can result in either I) complete removal of both C1q and C1r_2_s_2_ from the IgG-coated surface, or II) removal of only C1r_2_s_2_ while C1q remains stably bound to the IgG platform (**Fig. 4B**). Although C1-INH can inactivate C1r and C1s in solution too, it is important to note that we here only investigated the effect of C1-INH on C1 that is activated on an IgG-labelled surface.

Finally, our findings are also relevant for the correct interpretation of functional antibodies in research and diagnostics. C1q binding is often used as an important parameter to determine the presence of complement-fixing antibodies. For instance in transplantation, the serum of organ recipients is tested for the presence of donor-specific complement-binding anti-HLA antibodies (which could induce antibody-mediated rejection) by measuring binding of purified C1q ^80–83^. Based on our findings, direct C1q binding is an unreliable measure for downstream complement activation, as complement-binding IgGs that bind fully assembled C1, but not non-complexed C1q, are not detected. This also holds true for analyses of antibody-dependent complement activation in a human serum environment. For several antibodies, we observed that detection of surface-bound C1q is an unreliable read-out for IgG-dependent complement activation. This was most evident for wild-type IgG2 and IgG4-E430G, which could potently trigger C1-mediated complement activation (and phagocytosis), while we could hardly detect C1q on the surface. Although many studies have addressed the complement-binding and -activating properties of antibodies, surprisingly few have differentiated between C1q and C1. Our systematic investigation of C1q versus C1 binding to all IgG subclasses in both purified and human serum environments was key for the revelation of the importance of C1r_2_s_2_ in establishing stable C1q-IgG complexes.

## Acknowledgements

The authors kindly thank dr. Rob de Jong, dr. Annette Stemerding, dr. Lubka Roemenia and dr. Leendert Trouw and for scientific advice. We thank dr. Brandon Garcia for providing BBK32. This work was supported by an ERC Starting Grant (639209-ComBact, to S.H.M.R) and ERC Advanced Grant (233229-Coco, to P.G.), the Utrecht University Molecular Immunology Hub (UMI-Hub, ESTIMATE), and the Swiss National Science Foundation (31003A_176104, to DR). JP acknowledges support by the European Fund for Regional Development (EFRE, Regio 13), the Federal State of Upper Austria, and the Austrian Science Fund (FWF, P33958, P34164).

## Conflict of interest

AK, JAGS, PWHIP, KPMK, FJB, JS and SHMR are co-inventor on a patent describing antibody therapies against *S. aureus*.

---

**Supplemental Figure 1: Schematic overview of the classical complement pathway (A)** C1 binds surface-bound antibodies (either IgG or IgM) and, by cleaving C4 and C2, generates a surface-bound C3 convertase enzyme (C4b2b) that catalyzes the rapid deposition of C3b molecules onto the target surface. C3b evokes phagocytosis of the target cell, formation of C5 convertases and formation of the membrane attack complex (MAC). C1 binds to antibodies on the surface via the globular heads of subunit C1q. C1q binding requires oligomerization of surface-bound IgGs into ordered hexamers that are held together by non-covalent Fc-Fc contacts. **(B)** The C1 complex (C1qr_2_s_2_) consists of one recognition molecule C1q (green) and an associated protease heterotetramer consisting of C1r (pink) and C1s (purple) (C1r_2_s_2_). C1q consists of six collagen arms that each end in a globular domain (gC1q) at their C-terminus. Each gC1q domain can bind to the Fc tail of an IgG antibody.

**Supplemental Figure 2: IgG binding to *S. aureus* in serum** Binding of anti-WTA IgG (wild-type) to *S. aureus* Wood46 in 1% ΔIgGΔIgM serum. Data represent mean ± SD of three independent experiments.

**Supplemental Figure 3: C3b deposition on *S. aureus* in serum is C1-dependent**. C3b deposition on *S. aureus* Newman *Δspa/sbi* bacteria, labelled with either anti-WTA IgG1, IgG2 or IgG3 in 1% ΔIgGΔIgM serum in absence (‘control’) or presence of monoclonal anti-C1q antibody (blocking the gC1q) or BBK32 (blocking C1r-dependent cleavage of C4). Data represent mean ± SD of three independent experiments.

**Supplemental Figure 4: Verification of DNP-beads as model surface for antibody-dependent complement activation (A)** Schematic presentation of the DNP-bead model. Streptavidin-beads (with a diameter comparable to bacterial cells, 2.7 µm) are labelled with DNP-biotin, which is recognized by monoclonal anti-DNP IgGs. SA = streptavidin. **(B)** Streptavidin-beads were labelled with different concentrations of DNP-biotin and incubated with anti-DNP IgG1 and goat anti-human kappa-AlexaFluor647 in flow cytometry. Data represent mean ± SD of two independent experiments. **(C)** Binding of anti-DNP IgG subclasses to DNP-beads (1 µg/ml DNP) as determined by flow cytometry. Data represent mean ± SD of three independent experiments. **(D)** Binding of different forms of purified C1 to DNP-beads (1 µg/ml DNP) that were not labelled with IgG. C1q was detected by polyclonal anti-C1q antibodies and flow cytometry. Data represent mean ± SD of three independent experiments. **(E)** Binding of different forms of C1 to DNP-beads (1 µg/ml DNP) labelled with 20 nM IgG as detected by western blotting. After incubation with C1/C1q/C1-EDTA, beads were pelleted and supernatant was discarded. The bead pellet was boiled in SDS sample buffer for 5 minutes and then again pelleted. The supernatant, containing the previously surface-bound proteins, was next analysed by SDS-PAGE and western blotting with anti-C1q antibodies. The first two lanes show controls consisting of 3 nM purified C1q or C1. Representative blot of three experiments.

**Supplemental Figure 5: C1q binding to IgG-labelled *S. aureus* bacteria is similar to that in the DNP-bead model (A)** Binding of anti-WTA IgG (wild-type) to *S. aureus* Wood46 in RPMI buffer. **(B)** Binding of different forms of purified C1 to *S. aureus* Wood46 bacteria that were not labelled with IgG. ‘C1’ indicates the the fully assembled C1 complex (= C1qr_2_s_2_), ‘C1-EDTA’ sample consists of C1q, C1r and C1s, but the proteases are not attached to C1q (because 10 mM EDTA disrupts the Ca^2+^-dependent association between proteases and C1q). C1q was detected by polyclonal anti-C1q antibodies and flow cytometry. **(C)** Binding of purified C1 on *S. aureus* Wood46 bacteria labelled with 50 nM anti-WTA IgG, in the absence or presence of EDTA. C1q binding was detected by polyclonal anti-C1q antibodies and flow cytometry. **(A-C)** Data represent mean ± SD of three independent experiments.

**Supplemental Figure 6: IgG3 potently mediates C3b deposition on beads coated with lower DNP concentrations** C3b deposition in serum on beads loaded with a lower DNP concentration. Streptavidin-beads were loaded with 0.003 µg/ml biotinylated DNP antigen (∼300-fold lower than in previous experiments) and subsequently incubated with human monoclonal anti-DNP IgG (20 nM) and human ΔIgGΔIgM serum. Deposition of C3b molecules on the beads was determined by flow cytometry. Data represent geometric mean ± SD of two independent experiments.

**Supplemental Figure 7: Binding experiments by SPR (A)** Preparation of DNP and control surfaces on SPR sensor chip. A flat carboxymethyldextran surface was activated using NHS/EDC and aminated with ethylenediamine. A DNP construct with a short PEG spacer and amine-reactive NHS group was coupled to the active surface. Methoxy-PEG(4)-NHS was used to prepare a control surface with similar characteristics but missing the DNP antigen. **(B)** Binding of anti-DNP IgG1-4 (20 nM) to the DNP surface. All IgG isotypes associated to the active surface, providing a stable IgG baseline for subsequent C1 or C1q binding. Data show reference-subtracted signals on DNP surface. **(C)** Example of a typical experimental SPR injection sequence on DNP-labelled and control surfaces. IgG capturing and injection of C1 protein only induced detectable SPR signals on the DNP surface but not on the MeO control surface. Injection of glycine buffer (pH 2.0) led to a complete regeneration of the DNP surface. The example sequence shown here: IgG3-C1. **(D)** Binding of purified C1q and C1 to an SPR sensor chip subsequently coated with immobilized DNP and 4 nM anti-DNP IgG3 (5-fold lower than in **(B)**). C1q or C1 was injected for 60 seconds to allow association, after which the injection was stopped and dissociation was monitored. C1 dissociated slower than C1q alone, indicating that C1 binds more stable. **(B-D)** Data show one representative of two independent experiments. SPR responses were normalized to account for the molecular weight difference between C1 (766 kDa) and C1q (410 kDa). RU, reponse units.

**Supplemental Figure 8: Removal of C1r and C1s by C1-INH can result in dislodgement of C1q from surface-bound IgG (A-D)** Detection of C1r, C1s, C1-INH and C1q upon incubation of surface-bound C1 complexes with C1-INH. First, DNP-beads were labelled with anti-DNP IgG (20 nM IgG1 **(A-B)**, or IgG2, IgG3, IgG4 **(C-D)**) and C1 (1 nM), successively. Next, beads were incubated with C1-INH (200 nM) or a buffer control. Beads were pelleted and the supernatant was collected and incubated 1:1 (v/v) with SDS sample buffer (+DTT). **(A, C)** The presence of dissociated C1-INH–protease complexes in the supernatant samples was shown by western blotting with polyclonal anti-C1r, anti-C1s and anti-C1-INH. Red triangles indicate the complexes. Detection of C1-INH was only done for samples incubated with C1-INH. Controls shown (left-sided boxes) consist of purified C1 (3 nM), C1r (10 nM), C1s (10 nM) or C1-INH (10 nM). Representative blots of two experiments. *Explanation of bands:* Inhibition of C1r or C1s (associated to surface-bound C1q) occurs through covalent binding of C1-INH (110 kDa) to the C1r/C1s serine protease (SP) domain in the C1r/C1s light chain (LC), which separates from the C1r/C1s heavy chain under reducing conditions. Formation of inhibitor-protease complexes is thus visible by detection of C1-INH–C1r_LC_ (∼145 kDa) or C1-INH–C1s_LC_ (∼138 kDa) complexes, indicated by red triangles. **(B, D)** The dislodgement of C1q into the supernatant was determined by western blotting using polyclonal anti-C1q antibodies. Blue triangles indicate the presence of C1q chains A, B and C in the supernatant. Controls shown in the left-sided boxes consist of purified C1 (3 nM). Representative blot of two experiments.

**Supplemental Figure 9: IgG and C1q binding to DNP-beads coated with hexamer-enhanced IgG mutants (A)** Binding of anti-DNP IgG (wild-type, E430G and E345K mutants; 20 nM) to DNP-beads (1 µg/ml DNP) in PBS-TH (PBS with 0.05% (v/v) Tween and 0.5% human serum albumin). Unpaired *t* test (wild-type vs E430G or wild-type vs E345K) showed no significant differences (*P* > 0.05). **(B)** Binding of anti-DNP IgG (wild-type and E430G; 30 nM) to DNP-beads (1 µg/ml DNP) in 1% ΔIgGΔIgM serum. **(C)** Binding of C1-EDTA to DNP-beads (1 µg/ml DNP) labelled with anti-DNP IgG (wild-type, E430G or E345K mutants; 20 nM). ‘C1-EDTA’ sample consists of C1q, C1r and C1s, but the proteases are not attached to C1q (because 10 mM EDTA disrupts the Ca^2+^-dependent association between proteases and C1q). C1q binding was detected by polyclonal anti-C1q antibodies and flow cytometry. **(A-C)** Data represent geometric mean ± SD of three independent experiments.

**Supplemental Figure 10: Enhanced IgG oligomerization stabilizes C1q-IgG interactions on DNP-coated surfaces in SPR experiments** SPR experiments showing binding of purified C1 or C1q to sensor chips that are subsequently coated with immobilized DNP and with 20 nM anti-DNP IgG (either wild-type, E430G or E345K mutants). C1 or C1q was injected for 60 seconds to allow association, after which the injection was stopped and dissociation was monitored. Data show representative of two independent experiments. SPR responses were normalized to account for the molecular weight difference between C1 (766 kDa) and C1q (410 kDa). Data shown for wild-type IgG1-4 are identical to those shown in **Fig. 3A**. RU, reponse units.

**Supplemental Figure 11: Enhanced IgG oligomerization stabilizes C1q-IgG interactions on *S. aureus* bacteria (A)** Binding of anti-WTA IgG (wild-type and E430G mutant) to *S. aureus* Wood46 in RPMI buffer. Data shown for wild-type IgG1-4 are identical to those shown in **SFig. 5A. (B-C)** Binding of different forms of purified C1 on *S. aureus* Wood46 bacteria labelled with 50 nM anti-WTA IgG (either wild-type or E430G mutant). ‘C1-EDTA’ samples **(B)** consist of C1q, C1r and C1s, but the proteases are not attached to C1q (because 10 mM EDTA disrupts the Ca^2+^-dependent association between proteases and C1q). ‘C1’ **(C)** indicates the the fully assembled C1 complex (= C1qr_2_s_2_). C1q binding was detected by polyclonal anti-C1q antibodies and flow cytometry. Data shown for wild-type IgG1-4 are identical to those shown in **SFig. 5C. (A-C)** Data represent mean ± SD of three independent experiments.

**Supplemental Figure 12: Enhanced IgG oligomerization stabilizes C1q-IgG interactions on *S. aureus* bacteria in serum (A)** Binding of anti-WTA IgG (wild-type and E430G mutant) to *S. aureus* Wood46 in 1% ΔIgGΔIgM serum. Data represent mean ± SD of three independent experiments. Data shown for wild-type IgG1-4 are identical to those shown in **SFig. 2. (B)** Binding of C1q to anti-WTA IgG-labelled *S. aureus* Wood46 bacteria in 1% ΔIgGΔIgM serum as determined by flow cytometry. Data represent mean ± SD of three independent experiments. Data shown for wild-type IgG1-4 are identical to those shown in **Fig. 1B**.

**Supplemental Figure 13: Enhanced IgG oligomerization does not influence FcR-dependent phagocytosis of *S. aureus* bacteria by human neutrophils** Phagocytosis in the absence of complement. Phagocytosis of fluorescently labelled *S. aureus* Wood46 in RPMI buffer supplemented with anti-WTA IgG (wild-type or E430G mutant) and human neutrophils. Bacterial uptake was quantified by flow cytometry and displayed as the number of GFP-positive neutrophils relative to IgG1 wild-type. Data represent relative mean ± SD of three independent experiments. Phagocytosis data shown for wild-type IgG are identical to the data shown in **Fig. 6B**.

**Movie S1: HS-AFM of the IgG1-RGY – C1q complex in the presence of C1q in solution**. Initially detected C1q molecules subsequently dissociate from IgG hexamers. Image size: 400 x 400 nm^2^ (100 x 100 pixel^2^), 1 s /frame.

**Movie S2: HS-AFM of the IgG1-RGY – C1 complex in the absence of C1 in solution**. C1 molecules remain stably bound to IgG hexamers during HS-AFM imaging. Image size: 400 x 400 nm^2^ (100 x 100 pixel ^2^), 1 s /frame.

**Supplemental Table 1: Protein sequences used for antibody production** Variable and constant heavy and light chain protein sequences used for antbody production. The residues E345 and E430 are highlighted in grey.

## Methods

### Complement proteins

All complement proteins were obtained from Complement Technology Inc. (TX, USA): C1 (A098), C1q (A099), C1r (A102), C1s (A104) and C1-inhibitor (A140).

### Depletion of IgG and IgM from human serum

Human serum from 20 healthy donors was pooled and depleted of IgG and IgM as previously described ^33^. Informed consent was obtained from all subjects in accordance with the Declaration of Helsinki. Approval from the Medical Ethics Committee of the University Medical Center Utrecht was obtained (METC protocol 07-125/C approved on March 1, 2010). Briefly, IgG and IgM were separated from serum by affinity chromatography using HiTrap Protein G High Performance column (Cytiva, GE Healthcare) and CaptureSelect IgM Affinity Matrix (Thermofisher Scientific). Complement levels and complement activity were determined after depletion, using ELISA and classical/alternative pathway hemolytic assays respectively. Since C1q was partially co-depleted during the procedure, the ΔIgGΔIgM serum was reconstituted with C1q to physiological levels (70 µg/ml).

### Production of human monoclonal antibodies

Recombinant monoclonal anti-DNP (DNP-G2a2) IgG1 antibody with the triple RGY mutation _15_ used for HS-AFM experiments was obtained from Genmab (Utrecht, the Netherlands) ^16,84^. Other monoclonal human anti-DNP and anti-WTA IgG of the four IgG subclasses were produced recombinantly in human Expi293F cells (Life Technologies) as described before ^51^ with minor changes. The constant regions of heavy (HC) and light chain (LC) were cloned separately into pcDNA3.4 (ThermoFisher Scientific). Subsequently, gBlocks containing codon optimized variable (VH and VL) region sequences with an upstream KOZAK and HAVT20 signal peptide (Integrated DNA Technologies, IA, USA) were cloned upstream the HC and LC regions, respectively, using Gibson assembly (New England Biolabs). NheI and BsiWI were used as the 3’ cloning sites for VH and VL, respectively, in order to preserve the immunoglobulin heavy and kappa light chain amino acid sequence.

Sequences of IgG1, IgG2, IgG3, IgG4 variable and constant heavy and light chains were previously described ^51,85^ **(Supplemental Table 1)**. VH and VL sequences were derived from previously described anti-WTA GlcNAc-β-4497 (anti-WTA-4497; based on patent WO/2014/193722)^86^ and anti-DNP ^87^ **(Supplemental Table 1)**. Single point mutation E430G or E345K was introduced in the heavy chain expression vectors either by PCR-mediated site directed mutagenesis or by gene synthesis (IDT). Antibody amino acids are numbered according to EU nomenclature ^88^. After expression, IgG1, IgG2 and IgG4 antibodies were isolated from cell supernatant using a HiTrap protein A column (Cytiva, GE Healthcare). IgG3 antibodies were isolated using a HiTrap Protein G High Performance column (Cytiva, GE Healthcare). Antibodies were dialyzed overnight in PBS and filter-sterilized though 0.22 µm Spin-X filters. Antibodies were analyzed by size exclusion chromatography (SEC) (Cytiva, GE Healthcare) and separated for monomeric fraction in case aggregation levels were >5%. Antibody concentration was determined by measurement of the absorbance at 280 nm and antibodies were stored at -20°C.

### IgG and complement binding to DNP-coated beads

Magnetic Dynabeads M-270 Streptavidin (Invitrogen) (∼6-7 × 10^5^ beads per sample) were washed in PBS-TH (PBS, pH 7.4, 0.05% (v/v) Tween, 0.5% human serum albumin (HSA)) and incubated with 0.003 or 1 μg/ml biotinylated 2,4-dinitrophenol (DNP-PEG2-GSGSGSGK(Biotin)-NH2; 1186 Da; synthesized by Pepscan Therapeutics B.V., the Netherlands) in 0.1 ml/sample PBS-TH for 30 minutes at 4°C, shaking (±700 rpm). Next, DNP-coated beads were washed once in PBS-TH and incubated in 0.05 ml/sample PBS-TH with 20 nM anti-DNP IgG or a 3-fold serial dilution of anti-DNP IgG (starting from 90 nM) for 30 minutes at 4°C, shaking (±700 rpm). Subsequent incubations mentioned below were performed in 0.025 ml/sample PBS-TH for 30 minutes at 4°C, shaking (±700 rpm), unless otherwise stated. After each incubation, beads were washed three times with PBS-TH or VBS-TH (Veronal Buffered Saline pH 7.4, 0.25 mM MgCl_2_, 0.5 mM CaCl_2_, 0.05% (v/v) Tween, 0.5% HSA) dependent on the buffer used in the subsequent incubation step.

For detection of IgG binding, beads were incubated with 1 µg/ml goat anti-human kappa-AlexaFluor647 (Southern Biotech, 2060-31). Next, beads were washed and fixed in 0.15 ml/sample 1% paraformaldehyde in PBS-TH.

For C1q binding experiments, beads were incubated in 0.025 ml/sample VBS-TH with a 3-fold serial dilution (starting at 10 nM) of C1 or C1q for 30 minutes at 37°C, shaking (±700 rpm). To determine binding of ‘C1-EDTA’, C1 was incubated in VBS-THE (VBS-TH, 10 mM EDTA) instead. To study the effect of C1-INH and EDTA on C1q binding, beads were first incubated with 0.025 ml/sample 3 nM C1 and then with 0.025 ml/sample VBS-THE or 0.025 ml/sample 200 nM C1-INH in VBS-TH for 30 minutes at 37°C, shaking (±700 rpm). For C1q detection, beads were incubated with 5 μg/ml FITC-conjugated rabbit anti-human C1q (Dako, F0254).

To determine IgG, C1q and C3b deposition to beads in human serum, DNP-coated beads (∼6-7 × 10^5^ beads per sample) were incubated in 0.05 ml/sample VBS-TH with 20 nM anti-DNP IgG and a 3-fold serial dilution of ΔIgGΔIgM serum (starting from 10%) for 30 minutes at 37°C, shaking (±700 rpm). Otherwise, DNP-coated beads were incubated in VBS-TH with a 3-fold serial dilution of anti-DNP IgG (starting from 90 nM) and 1% ΔIgGΔIgM serum for 30 minutes at 37°C, shaking (±700 rpm).

For IgG, C1q and C3b detection, beads were washed and incubated in 0.025 ml/sample PBS-TH with a detection antibody for either IgG, C1q or C3b (1:300 FITC-conjugated goat anti-C3 IgG, De Beer Medicals, 1100). After incubation with detection antibodies, bead samples were washed and fixed in 0.15 ml/sample 1% paraformaldehyde in PBS-TH and binding of IgG, C1q or C3b to the beads was determined by flow cytometry (BD FACSVerse). Data were analyzed based on single bead population using FlowJo software and presented as Fluorescence Intensity (FI) means ± SD or geometric means ± SD of at least three independent experiments.

### Western blotting

To determine binding of different forms of C1 to DNP-beads by western blotting, the amount of beads (and incubation volumes) were scaled up 4-fold (∼24-28 × 10^5^ beads/sample). Magnetic Dynabeads M-270 were successively incubated with 1 µg/ml DNP-biotin, 20 nM anti-DNP IgG and 3 nM C1q/C1/C1-EDTA, as described above. Next, beads were washed in VBS-TH and boiled in 50 µL sample buffer (0.5 M Tris-HCl, 2% (w/v) SDS, 20% (v/v) glycerol, 0.1% (w/v) bromophenol blue, pH 7) with 25 mg/ml dithiothreitol (DTT) for 5 minutes at 95°C. Beads were pelleted and 10 µL of the sample supernatant was run on a 10% SDS-PAGE gel and subsequently transferred to a PVDF membrane (Trans-Blot Turbo Transfer System, Bio-Rad). Membranes were blocked with 4% dried skim milk (ELK, Campina) in PBS-T (PBS with 0.05% Tween) for 1 hour at 37°C. To detect C1q, membranes were incubated for 45 minutes in 1% dried skim milk in PBS-T with rabbit anti-human C1q (1:300, Dako, A0136) and HRP-conjugated goat anti-rabbit IgG (1:10.000, Southern Biotech, 4030-05). Blots were washed between the successive antibody incubations.

To determine the removal of C1r and C1s from bead-bound C1 by C1-INH by western blotting, the amount of beads (and incubation volumes) were scaled up 3-fold (∼18-21 × 10^5^ beads/sample). Magnetic Dynabeads M-270 were successively incubated with 1 µg/ml DNP-biotin and 20 nM anti-DNP IgG, as described above. Next, beads were incubated with 1 nM C1 in VBS-T (Veronal Buffered Saline pH 7.4, 0.25 mM MgCl_2_, 0.5 mM CaCl_2_, 0.05% (v/v) Tween) for 30 minutes at 37°C, shaking. After washing, beads were incubated with 200 nM C1-INH or a VBS-T buffer control for 30 minutes at 37°C, shaking. Next, beads were pelleted and supernatant was collected and mixed 1:1 with 2x sample buffer containing 50 mg/ml DTT for 5 minutes at 95°C. Next, 10 µL samples were run on a 10% SDS-PAGE gel, transferred to a PVDF membrane and membranes were blocked, as described above. To detect C1r, C1s, C1-INH or C1q, the membrane was incubated in PBS-T with 1% ELK for 45 minutes with either goat anti-human C1r (1:300, R&D Systems, AF1807), sheep anti-human C1s (1:300, R&D Systems, AF2060), rabbit anti-C1-INH (1 µg/mL, Sino Biologicals, 10995-R018) or rabbit anti-human C1q (1:300), respectively. After washing, membranes were incubated 1:10.000 for 45 minutes with HRP-conjugated antibodies, either donkey anti-goat IgG (Southern Biotech, 6425-05), rabbit anti-sheep IgG (Southern Biotech, 6425-05) or goat anti-rabbit IgG. Membranes were developed using Enhanced Chemiluminesence (ECL, Cytiva, GE Healthcare).

### IgG and complement binding to *S. aureus*

Depending on the experiment, *S. aureus* Wood46 (with low expression of staphylococcal protein A (Spa)) or *S. aureus* Newman *Δspa/sbi* (knock-out of Protein A and the second immunoglobulin-binding protein (Sbi)) ^89^ was or was not fluorescently labelled by transformation with a GFP-expressing plasmid ^90^. Bacteria were grown until log phase in Todd Hewitt Broth medium, washed and frozen at -20°C until use. To prepare samples, bacteria were thawed and diluted to OD=0.1 in RPMI-H (RPMI, 0.05% HSA). All subsequent incubations mentioned below were performed in RPMI-H at 4°C, shaking (±700 rpm), unless stated otherwise. After each incubation, bacteria were washed once in RPMI-H by centrifugation. For detection of IgG binding, 0.025 ml bacteria was incubated with 0.025 ml 3-fold serial dilution anti-WTA IgG (starting from 90 nM, final concentration). Next, bacteria were incubated in 0.025 ml/sample RPMI-H with 1 µg/ml goat anti-human kappa-AlexaFluor647. For C1q binding experiments, 0.025 ml bacteria was incubated with 0.025 ml 50 nM anti-WTA IgG (final concentration). Bacteria were next incubated in 0.025 ml/sample RPMI-H with a 3-fold serial dilution of C1 or C1q (starting at 30 nM) for 30 minutes at 37°C, shaking (±700 rpm). To determine binding of ‘C1-EDTA’, C1 was incubated in RPMI-HE (RPMI-H, 10 Mm EDTA) instead. For C1q detection, bacteria were incubated with 5 μg/ml FITC-conjugated rabbit anti-human C1q.

To determine IgG, C1q and C3b binding to bacteria in serum, 0.050 ml bacteria were mixed with 0.025 ml 3-fold serial dilution anti-WTA IgG (starting from 90 nM, final concentration) and 0.025 ml 1% (IgG or C1q detection) or 5% (C3b detection) ΔIgGΔIgM serum (final concentration) and incubated 30 minutes at 37°C, shaking (±700 rpm). For IgG, C1q or C3b detection bacteria were next incubated with detection antibody for IgG, C1q or C3b (1:300 FITC-conjugated goat anti-C3 IgG).

To determine C3b deposition on bacteria in serum in the presence of C1-blocking molecules, 0.020 ml bacteria were mixed with 0.030 ml RPMI-H with a 3-fold serial dilution anti-WTA IgG (starting from 67 nM, final concentration), 1% ΔIgGΔIgM serum (final concentration) and either monoclonal anti-C1q antibody (HB8327 a4b11 ^46^, ATCC, 10 µg/ml final concentration) or BBK32 (kindly provided by Dr. Brandon Garcia, 10 µg/ml final concentration). Bacteria were incubated 20 minutes at 37°C, shaking (±700 rpm). For C3b detection bacteria were next incubated with FITC-conjugated goat anti-C3 IgG (1:300).

After incubation with detection antibodies, bacteria were washed and fixed in 0.15 ml/sample 1% paraformaldehyde in RPMI-H and analyzed using flow cytometry (BD FACSVerse). Data were analyzed by FlowJo software and presented as Fluorescence Intensity (FI) means ± SD or geometric means ± SD of at least three independent experiments.

### Phagocytosis

Human neutrophils were isolated from blood of healthy donors by the Ficoll-Histopaque gradient method ^91^. Phagocytosis was performed in a round-bottom 96-well plate. GFP-labelled Wood46 bacteria (20 µL of 3.75 × 10^7^ cells/ml) were mixed with 2-fold serial dilutions of anti-WTA IgG in RPMI-H or 1% ΔIgGΔIgM serum (20 µL volume) for 15 minutes at 37°C for opsonization. Subsequently, neutrophils (10 µL of 7.5 × 10^6^ cells/ml) were added giving a 10:1 bacteria to cell ratio and incubated for 15 minutes at 37°C on a shaker (750 rpm) in a final volume of 50 µL. The reaction was stopped with 1% ice-cold paraformaldehyde and neutrophil-associated fluorescent bacteria were analyzed by flow cytometry by scatter gating on neutrophils. Phagocytosis was defined by the percentage of cells with a positive fluorescent signal (% GFP-positive cells) of all neutrophils representing the overall phagocytosis efficacy.

### Surface plasmon resonance studies

All surface plasmon resonance experiments were performed at 25°C using a Biacore T200 instrument (Cytiva, GE Healthcare) equipped with a flat carboxymethyldextran sensor chip (CMDP; Xantec). For DNP surface preparations, flow cells were activated for 7 minutes with a 1:1 mixture of 0.1 M N-hydroxysuccinimide (NHS) and 0.1 M (3-(N,N-dimethylamino)propyl-N-ethylcarbodiimide) at a flow rate of 10 µL/min, followed by a 7 minute injection of 0.1 M ethylenediamine in 0.1 M sodium borate pH 8.5. DNP-NH-PEG(4)-NHS (Iris Biotech) was reacted at a concentration of 5 mM in PBS-T with the aminated surface to reach a coupling density of 250 reponse units (RU). As a reference surface, MeO-PEG(4)-NHS (20 mM) was coupled to a density of 200 RU as described for the DNP surface. Binding analyses were performed in HBST^++^ (10 mM HEPES, 150 mM NaCl, 0.5 mM CaCl_2_, 0.25 mM MgCl_2_, 0.005% Tween 20, pH 7.4) at a flow rate of 10 µL/min if not stated otherwise. Anti-DNP antibodies at concentrations of 4 nM and 20 nM were prepared in HBST^++^ and injected for 120 seconds over the DNP and control surfaces. Subsequently, C1 or C1q (20 nM) were injected for 60 seconds to associate on top of the antibodies and dissociation was observed for 300 seconds. The surfaces were regenerated with a 60 second injection of 10 mM glycine pH 2.0 at a flow rate of 20 µL/min. Data were collected at a rate of 1 Hz and analyzed with Scrubber (version 2.0c; BioLogic). Reference signals from the control surface (MeO) were subtracted from the DNP surface signals. The responses were normalized for analyte size contribution by dividing the data points of each analyte by the molecular weight of C1 (766 kDa) or C1q (410 kDa), respectively, and multiplying them by 100. The data were exported to Prism (version 8.1.2.; Graphpad).

### DNP-labelled liposomes

DNP-labelled liposomes consisting of 1,2-dipalmitoyl-sn-glycero-3-phosphocholine (DPPC), 1,2-dipalmitoyl-sn-glycero-3-phosphoethanolamine (DPPE) and 1,2-dipalmitoyl-sn-glycero-3-phosphoethanolamine-N-[6-[(2,4-dinitrophenyl)amino]hexanoyl] (DNP-cap-DPPE) were used to generate supported lipid bilayers (SLBs) on mica and SiO_2_ substrates. The lipids were purchased from Avanti Polar Lipids, mixed at a ratio of DPPC:DPPE:DNP-cap-DPPE = 90:5:5 (molar ratio), and dissolved in a 2:1 mixture of chloroform and methanol. After the solvents were rotary-evaporated for 30 minutes, the lipids were again dissolved in chloroform, which was then rotary-evaporated for 30 minutes. Drying was completed at a high vacuum pump for 2 hours. The lipids were dissolved in 500 µL Milli-Q H_2_O while immersed in a water bath at 60°C, flooded with argon, and sonicated for 3 minutes at 60°C to create small unilamellar vesicles. These were diluted to 2 mg/ml in buffer #1 (10 mM HEPES, 150 mM NaCl, 2 mM CaCl_2_, pH 7.4) and frozen for storage using liquid N_2_.

### High-speed atomic force microscopy

HS-AFM ^92,93^ (RIBM, Japan) was conducted in tapping mode at RT in buffer, with free amplitudes of 1.5 -2.5 nm and amplitude set points larger than 90%. Silicon nitride cantilevers with electron-beam deposited tips (USC-F1.2-k0.15, Nanoworld AG), nominal spring constants of 0.15 N m^-1^, resonance frequencies around 500 kHz, and a quality factor of approx. 2 in liquids were used. Imaging was performed in buffer #1. All IgGs were diluted and incubated in the same buffer.

DNP-labelled supported lipid bilayers for HS-AFM were prepared on muscovite mica. The liposomes were incubated on the freshly cleaved surface (500 µg/ml in buffer #1), placed in a humidity chamber to prevent evaporation, and heated to 60 °C for 30 minutes. Then the temperature was gradually cooled down to RT within 30 minutes, followed by exchanging the solution with buffer #1. After 10 minutes of equilibration at RT and 15 more buffer exchanges, the SLB was ready for imaging. In order to passivate any exposed mica, SLBs were incubated with 333 nM IgG1-b12 (irrelevant human IgG1 control antibody against HIV-1 gp120) ^94^ for 10 minutes before the molecules of interest were added. Anti-DNP IgG1-RGY (20 µg/ml) was incubated for 5 minutes followed by several buffer exchanges to remove umbound IgGs from solution. After that, C1q (20 µg/ml) or C1 (15 µg/ml) was added for 10 minutes, followed by another buffer exchange. In case of C1q, imaging was performed in the presence of C1q in soluiton to enable rebinding during the experiment.

### Statistical analysis

Statistical analysis was performed with Prism software (version 8.0.1; GraphPad). All data are presented as means ± SD or geometric means ± SD from three independent experiments unless otherwise stated.

